# A lineage-specific heat-induced feedback loop controls HrcA to promote chlamydial fitness under stress

**DOI:** 10.1101/2025.05.30.657042

**Authors:** Yehong Huang, Yuxuan Wang, Matthew Pan, Danny Wan, Lingling Wang, Joseph D. Fondell, Xiang Wu, Guangming Zhong, Huizhou Fan

## Abstract

Bacterial stress responses rely on transcriptional regulation by sigma factors and repressors. In *Chlamydia*, which lacks a dedicated heat shock sigma factor, repressor HrcA limits chaperone gene expression under non-stressed conditions. While HrcA function may be enhanced by the chaperone GroESL in a positive feedback loop, here we identify a heat-induced negative feedback loop mediated by HagF, a protein unique to Chlamydiota and conserved across *Chlamydia* species. We show that HrcA represses *hagF*, but heat shock induces *hagF* expression, leading to HagF accumulation and binding to HrcA. Structural modeling and pulldown assays indicate that HagF blocks HrcA dimerization, impairing DNA binding. This relieves repression of the *hrcA-grpE-dnaK* operon and promotes secondary differentiation in the developmental cycle, enabling infectious elementary bodies to form under heat stress. Our findings reveal dual feedback regulation that tunes chaperone gene expression and illustrate how transcriptional repression can be modulated in a minimal-genome pathogen.

## Introduction

The ability of cells to respond to environmental stress is fundamental to survival, with the heat shock response being among the most studied mechanisms of cellular defense ^1-5^. This universally conserved response equips organisms to withstand and recover from the detrimental effects of elevated temperatures and other stressors through transcriptomic reprogramming ^1,3-5^. In many bacteria, the upregulation of stress-responsive genes is mediated by specialized sigma factors within the RNA polymerase holoenzyme that recognize heat shock promoters ^1,3-5^. Additionally, specific genes can be upregulated following the dissociation of heat-inducible transcriptional repressors from their target promoters ^1,3-5^. HrcA is the most widely distributed transcriptional repressor among bacteria.

Members of the phylum Chlamydiota, including the human pathogen *Chlamydia trachomatis*, are obligate intracellular bacteria with a distinctive developmental cycle that alternates between the infectious elementary body (EB) and the intracellular, replicative reticulate body (RB) ^6-8^. Under extreme stress, RBs cease division but may continue genome replication, resulting in enlarged, non-dividing forms known as aberrant bodies. This condition, termed chlamydial persistence, is reversible upon return to favorable conditions, when aberrant bodies convert back to dividing RBs and resume progeny EB formation ^9-14^. In response to milder stress, some RBs may enter persistence while others continue to replicate slowly, eventually allowing EB production. The ability to sustain or recover EB formation under stress is essential for *Chlamydia* to maintain infectivity and transmission and represents a critical determinant of its pathogenesis^11,12,14,15^.

Despite possessing one of the smallest bacterial genomes and lacking a designated heat shock sigma factor, *Chlamydia* is capable of mounting a robust heat shock response characterized by extensive transcriptomic reprogramming ^5^. This suggests expanded roles for other stress regulators. Indeed, heat-induced expression changes in numerous genes have been linked to alterations in the expression and activities of the remaining sigma factors (i.e., σ66, σ28, and σ54) ^5^, thought mainly to control chlamydial growth and development.

HrcA is the only known heat-inducible transcriptional repressor in *Chlamydia*. Similar to its role in other bacteria ^16-19^, HrcA directly regulates two operons in *C. trachomatis* encoding the chaperones DnaK, GrpE, GroES, and GroEL, as well as HrcA itself ^20,21^. Under non-stressed conditions, HrcA binds CIRCE (controlling inverted repeat of chaperone expression) elements upstream of these operons and represses their transcription ^16-21^. This repression is believed to be stabilized by GroEL or the GroESL complex, forming a positive feedback loop that helps maintain HrcA activity in the absence of stress ^22-24^. During heat shock, HrcA becomes destabilized, leading to derepression of chaperone genes. Here, we identify *ctl0271*, a gene repressed by HrcA and induced by heat shock, as encoding a novel feedback inhibitor of HrcA. We term this gene *hagF* (heat-activated HrcA-repressed gene F). HagF binds to HrcA and interferes with its dimerization, thereby amplifying derepression of heat shock genes. This combination of positive feedback through GroESL under non-stress conditions and negative feedback through HagF during heat stress provides a temperature-dependent mechanism that fine-tunes chaperone expression. HagF is conserved across Chlamydiota but absent from other bacterial lineages and is essential for efficient EB formation under heat stress. These findings position HagF as a lineage-specific regulator that couples transcriptional derepression with developmental progression, thereby enhancing *Chlamydia* fitness and promoting dissemination from hosts mounting responses that restrict chlamydial growth.

## RESULTS

### *ctl0271 (hagF)* is a novel and the 6^th^ heat-inducible gene repressed by HrcA

Currently, HrcA is known to regulate the expression of only five genes by targeting two promoters in *Chlamydia* (Fig. 1A, B) ^20,21^. We suspected *ctl0271* as a novel HrcA target gene because *ctl0271* is strongly upregulated by heat shock ^5^, and its promoter region contains a putative CIRCE element (Fig. 1C). This putative CIRCE element is five nucleotides upstream of the -35 core promoter element, same as the CIRCE elements in KP1 and EP. The sequence of the putative CIRCE element of *ctl0271* deviates from that of the consensus sequence of bacterial CIRCE elements ^25^ by only two nucleotides, which is one more than that of the CIRCE element of KP1, but two fewer than that of EP, although the two established CIRCE elements both form stronger stem regions near the nine-nucleotide loop. However, the putative *ctl0271* CIRCE element possesses both higher sequence conservation and a more stable stem region, compared to an HrcA target in *Sinorhizobium* (formerly *Rhizobium*) *meliloti* ^26^ (Fig. 1C). These findings suggest that chlamydial HrcA potentially regulates the expression of *ctl0271*, in addition to the *hrcA-grpE-dnaK* and *groES-groEL* operons.

**Fig. 1.**
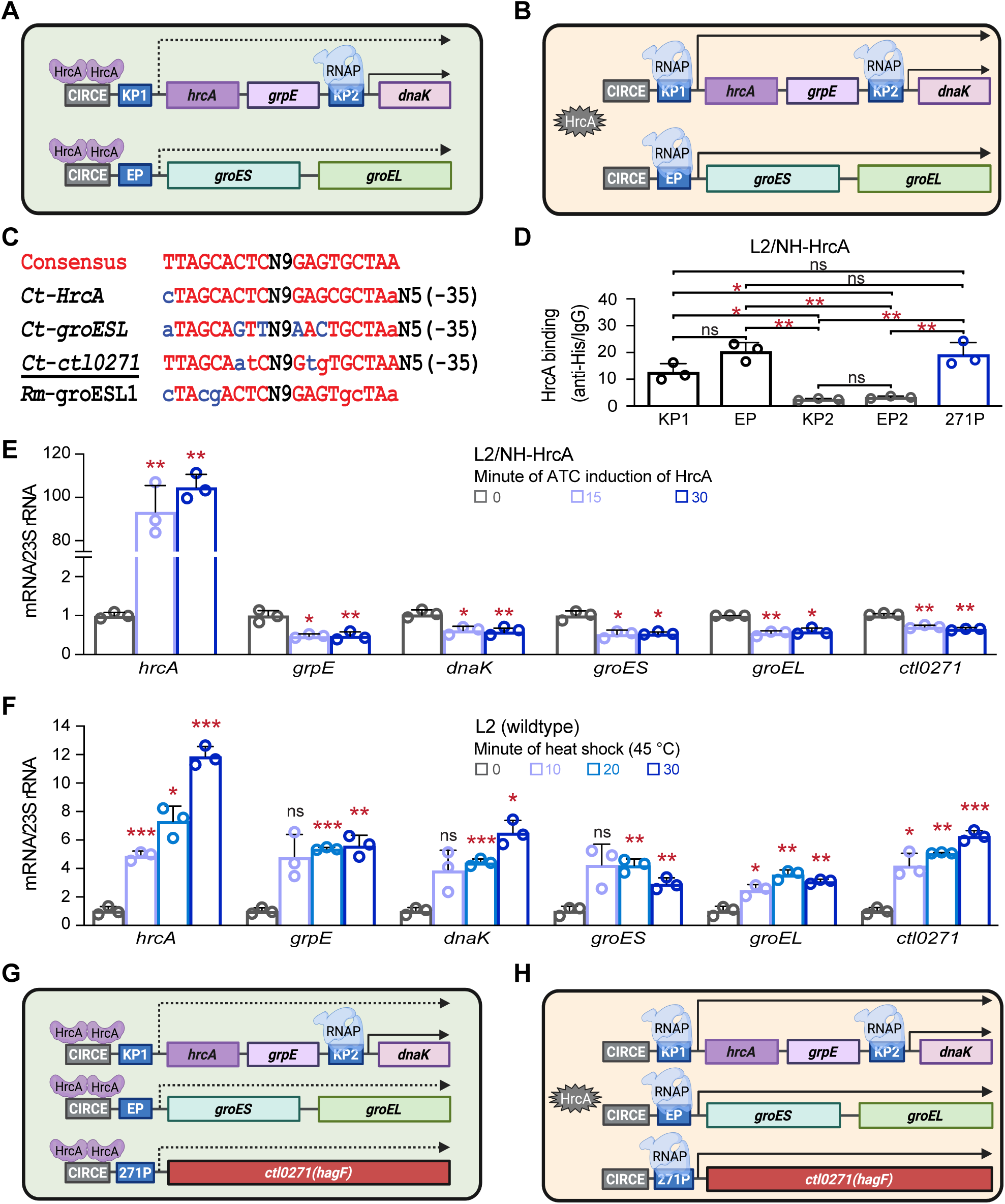
*ctl0271 (hagF)* is the 6th heat-inducible HrcA-repressed gene. (A) Under non-stressed conditions, HrcA binds to CIRCE elements and represses transcription from the KP1 and EP promoters. (B) Heat shock disrupts HrcA repression, allowing RNA polymerase (RNAP) to transcribe the *hrcA-grpE-dnaK* and *groES-groEL* operons. Arrows denote the extent of transcription. Note that *dnaK* is regulated by both the operon promoter and its own internal promoter (KP2). (C) Consensus and putative CIRCE sequences upstream of indicated heat shock genes in *C. trachomatis (Ct)* and *Rhizobium meliloti (Rm)*. Blue letters indicate nucleotide deviations from the consensus; lowercase letters denote mismatches between inverted repeats. The-35 core promoter elements for *C. trachomatis* genes are shown. (D) HrcA binds the *ctl0271* promoter in chlamydiae. ChIP-qPCR was performed in biological triplicate using L2/NH-HrcA cultures treated with 10 nM ATC for 30 min at 37 °C. Anti-His antibody enrichment of the *ctl0271* promoter was comparable to that of known HrcA targets (*groESL* and *dnaK* P1), and significantly higher than enrichment at non-target promoters KP2 and E2P (*groEL2* promoter) ^22,45^. (E) Expression of *ctl0271*, *dnaK*, and *groEL* was reduced upon ATC-induced NH-HrcA overexpression in L2/NH-HrcA, as determined by qRT-PCR. (F) *ctl0271* expression is induced in wild-type L2 cultured at 45 °C, as measured by qRT-PCR. (D–F) Data represent the mean ± SD of biological triplicates. (G, H) Based on results from panels B–F, the updated HrcA regulon includes *ctl0271* (renamed *hagF*) as its sixth target gene. (A, B, G, H) These panels were generated using a paid subscription to bioRender.

We conducted ChIP-qPCR analysis to confirm that HrcA binds to the *ctl0271* promoter in *C. trachomatis* L2 (L2) transformed with a plasmid carrying a recombinant gene expressing His-tagged HrcA (L2/NH-HrcA) cultured with ATC. Although the anti-HIS antibody enriched HrcA nontarget promoters *dnaK-*P2 (KP2) and *groEL2-*P (E2P) over the control IgG by 2.5- and 3.2-fold, respectively, anti-HIS enriched KP1 and EP by 12.6- and 20.4-fold, respectively (Fig. 1D). Significantly, anti-HIS also enriched the *ctl0271* promoter (271P) by 19.1-fold (Fig. 1D). This results revealed that NH-HrcA binds to 271P in *C. trachomatis*.

Based on the results of sequence analysis (Fig. 1C) and ChIP-qPCR (Fig. 1D) analyses and the role of HrcA as a transcriptional repressor (Fig. 1A, B), we hypothesized that *HrcA represses ctl0271 expression*. To test this hypothesis, we first performed quantitative real-time reverse transcription PCR (qRT-PCR) analysis in L2/NH-HrcA. Supporting this hypothesis, induction of NH-HrcA expression with ATC at 37 °C for 15 and 30 min resulted in statistically significant -1.4 and -1.5 fold changes in *ctl0271* transcripts (Fig. 1E), respectively, compared with control non-induction. These decreases are comparable to expression decreases in *grpE, danK, groES,* and *groEL* (Fig. 1E). We next determined the effect of heat shock, which disables HrcA to bind its target promoters on *ctl0271* expression in cultures of wildtype L2. Incubation at 45 ℃ for 10, 20, and 30 minutes resulted in 4.2, 5.1, and 6.3-fold increases in *ctl0271* expression, respectively (Fig. 1F), compared to incubation at 37 ℃. These increases are also comparable to those in *hrcA, grpE, dnaK, groES,* and *groEL.* The multiple-fold expression increases in response to heat shock and only modest expression reduction in response to massive *hrcA* overexpression at 37 ℃ are consistent with the notion that the 271P, like KP1 and EP, is mostly repressed by HrcA in chlamydiae cultured at 37 ℃, and strongly de-repressed by heat shock.

Taken together, findings presented in Fig. 1C-F indicate that *ctl0271* is a novel and the 6^th^ HrcA-repressed heat-activated gene in *C. trachomatis* (Fig. 1F). Therefore, we renamed *ctl0271 hagF*. Notably, HagF is conserved in all *Chlamydia* species and *Chlamydia-*like organisms (Fig. S1). Although BlastP analysis using *C. trachomatis* HagF as a query revealed that one non-chlamydia bacterium, *Bacteroides fragilis* S6L5, might carry *hagF* (Fig. S2A). Further analysis showed that the predicted *B. fragilis* B6L5 protein sequence is identical to that of the HagF ortholog in *C. gallinacea* (Fig. S2B), suggesting that the *B. fragilis* genomic DNA sample in that sequence analysis was likely contaminated with *C. gallinacea* genomic DNA. The finding of HagF as an exclusively *Chlamydia*-specific protein suggests that HagF evolved to play a critical role in chlamydial physiology.

### HagF promotes EB formation during heat shock

In addition to regulating stress responses, some HrcA target genes are known to be essential for bacterial growth under non-stress conditions ^27,28^. To probe the physiological role of HagF, we employed deactivated CRISPR-associated protein 12 (dC12) ^29^ to silence *hagF* expression in *C. trachomatis*. We derived three L2 transformants, each expressing dC12 in an ATC-inducible manner and a dC12 guide RNA (gRNA) constitutively. While gRNA g1 and g2 target different regions of *hagF,* ntg targets a synthetic gene not found in *Chlamydia* (see Materials and Methods). As expected, in the transformant L2/dC12-ntg, induction of dC12 expression with ATC did not affect the expression level of *hagF* (Fig. 2A). In comparison, in transformants L2/hagF*-*dC12-g1 and L2/hagF*-*dC12-g2, induction of deactivated Cas12 expression (Fig. 2A, left) resulted in -5.4 and -3.1 fold changes, respectively, in the level of *hagF* transcripts (Fig. 2A), suggestive of effective silencing of *hagF* expression by both gRNAs.

**Fig. 2.**
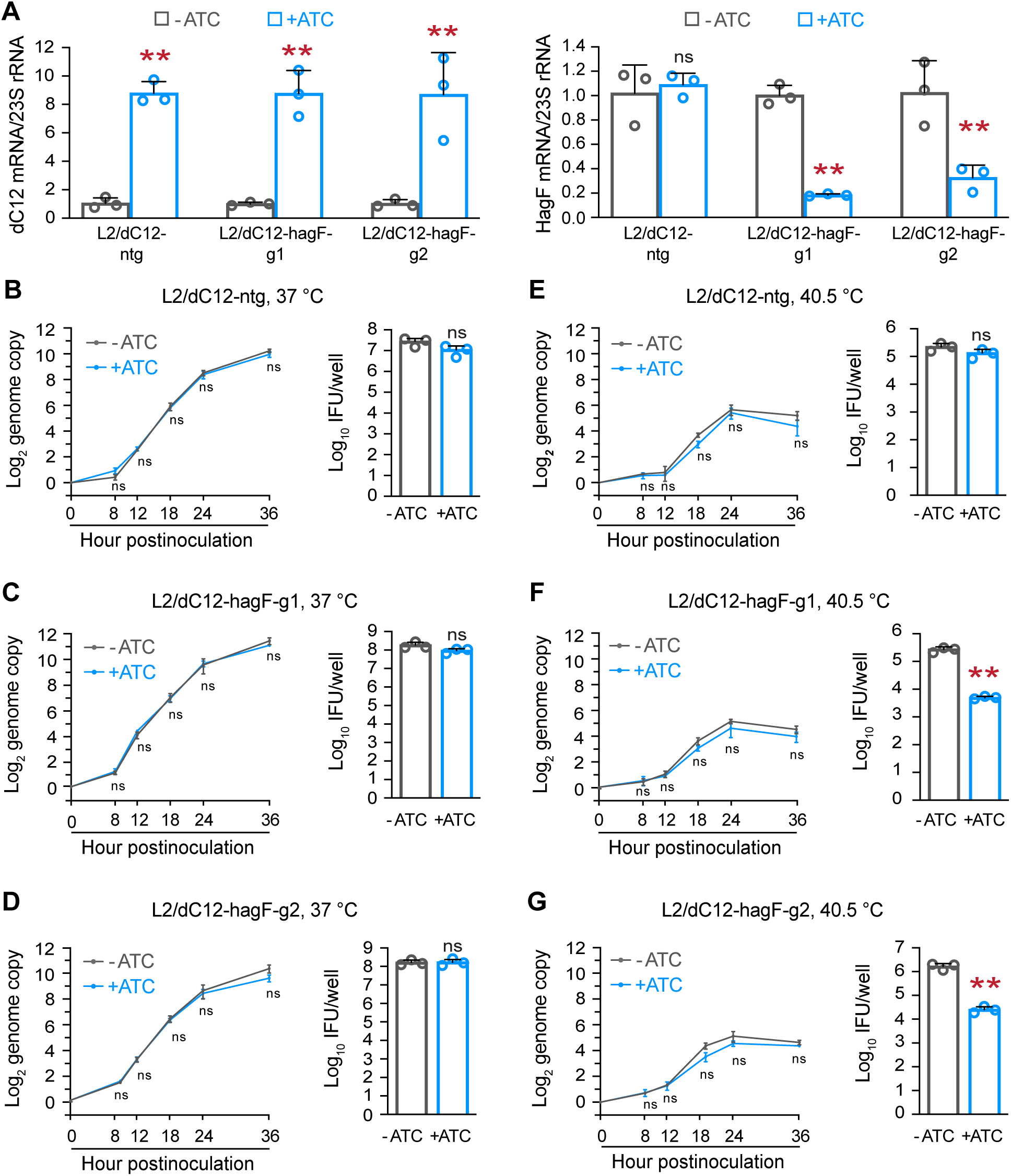
*hagF* facilitates EB formation during heat shock. (A) Effective silencing of *hagF* in *C. trachomatis* using deactivated CRISPR-associated protein 12 (dC12). L2/dC12-ntg expresses an ATC-inducible *dC12* and a non-targeting guide RNA. L2/dC12-hagF-g1 and L2/dC12-hagF-g2 express *dC12* and a guide RNA targeting *hagF* (g1 or g2). The left panel shows elevated *dC12* transcript levels upon ATC induction in all three transformants. The right panel shows reduced *hagF* expression in ATC-treated L2/dC12-hagF-g1 and L2/dC12-hagF-g2 but not in L2/dC12-ntg. RNA levels were quantified by qRT-PCR. (B) ATC-induced *dC12* expression has no effect on genome replication (left) or EB formation (right) in 37 °C cultures of L2/dC12-ntg. (C, D) ATC-induced *hagF* silencing has no detectable effect on genome replication (left) or EB formation (right) in 37 °C cultures of L2/dC12-hagF-g1 (C) and L2/dC12-hagF-g2 (D). (E) ATC-induced *dC12* expression has no effect on genome replication (left) or EB formation (right) in 40.5 °C cultures of L2/dC12-ntg. (F, G) ATC-induced *hagF* silencing does not affect genome replication (left) but causes a substantial reduction in EB formation (right) in 40.5 °C cultures of L2/dC12-hagF-g1 (F) and L2/dC12-hagF-g2 (G). (B-G) ATC was added at 0 hpi. Genome copy number was quantified at the indicated times. EB yields were measured at 30 hpi for 37 °C cultures and at 40 hpi for 40.5 °C cultures.

When L2/dC12-ntg was cultured at either 37 ℃, the genome replication kinetics (Fig. 2B) in ATC-containing and ATC-free cultures was indistinguishable, as was the EB yield, suggesting that dC12 expression does not nonspecifically inhibit chlamydial RB replication or progeny EB formation. When L2/dC12-hagF-g1 and L2/dC12-hagF-g2 were cultured at 37 ℃, ATC-induced *hagF* silencing through either guide RNA did not affect the genome replication kinetics or EB formation (Fig. 2C and D), suggesting that *hagF* is not an essential gene for chlamydial growth and development under normal culture conditions.

We next assessed the effects of HagF silencing in L2/dC12-ntg, L2/dC12-hagF-g1, and L2/dC12-hagF-g2 at 40.5 ℃ (Fig. 2E-G). As expected, the genome replication kinetics occurs at a slower kinetics, and EB yields are lower in the 40.5 ℃ cultures (Fig. 2E-G), compared to the 37℃ cultures (Fig. 2B-D). Similar to 37 ℃ cultures, ATC-induced dC12 expression in L2/dC12-ntg did not affect either genome replication or EB formation (Fig. 2E). ATC-induced *hagF* silencing in L2/dC12-hagF-g1 and L2/dC12-hagF-g2 did not affect their genome kinetics either but reduced the EB yield by nearly 70-fold (Fig. 2F, G). These results suggest that HagF promotes EB formation under heat stress.

### HagF interacts with HrcA to block its transcriptional repression activity

To determine how HagF facilitates progeny EB formation under heat stress, we investigated its localization, which has been controversial in the literature ^30,31^. Although *Yersinia enterocolitica* type III secretion system was capable of secreting a β-lactamase fused to the first 20 amino acids of HagF, it failed to secrete the full-length HagF ^30^. However, another study detected the secretion of HagF fused to reporter enzymes from transformed chlamydiae to host cells ^31^. We employed an ATC-inducible system to express and localize HagF carrying either an N- or C-terminal HIS tag (i.e., NH-HagF and CH-HagF) to eliminate potential interference with secretion by the tag ^32-38^. Following ATC induction for 6 h, both NH-HagF (Fig. 3A) and CH-HagF (Fig. 3B) displayed granular signals superimposable with the signals of GrgA, a chlamydial cytoplasmic protein (transcription factor) ^39,40^. These results indicate that HagF is also a cytoplasmic protein.

**Fig. 3.**
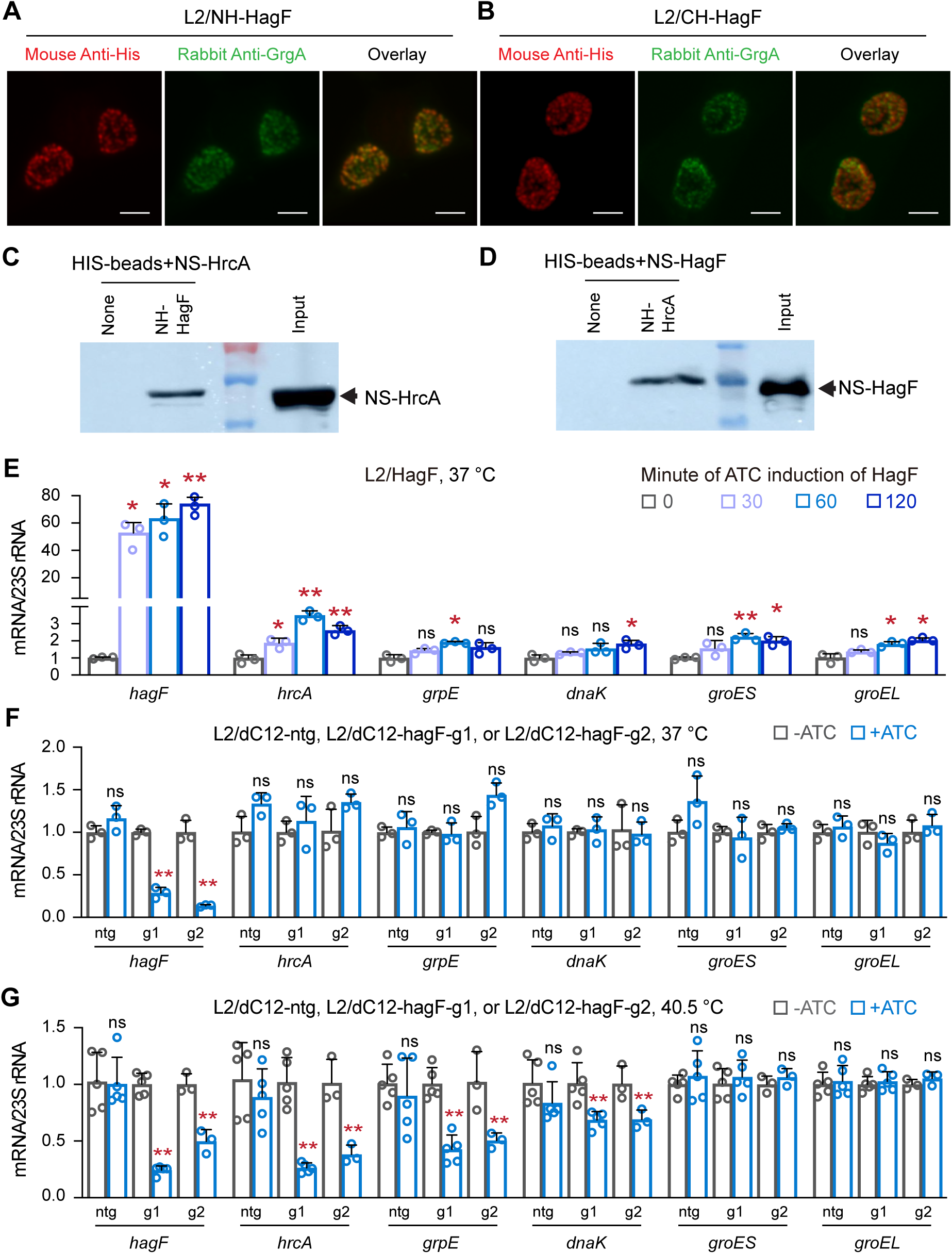
HagF binds to HrcA to inhibit its transcriptional repression activity. (A, B) HagF localizes inside chlamydial cells. HeLa 229 cells were infected with L2/NH-HagF (A) or L2/CH- HagF (B), fixed at 20 hpi, and subjected to immunostaining. NH-HagF and CH-HagF were detected using a mouse anti-His antibody and TRITC-conjugated goat anti-mouse IgG. GrgA, a cytoplasmic chlamydial transcription factor, was detected using a rabbit anti-GrgA antibody and FITC-conjugated goat anti-rabbit IgG. (C, D) HagF physically interacts with HrcA. In panel C, purified NS-HrcA (Strep-tagged HrcA) was incubated with HIS beads in the presence or absence of NH-HagF. In panel D, purified NS-HagF was incubated with HIS beads in the presence or absence of NH-HrcA. Following washes, bead-bound proteins were resolved by SDS-PAGE and detected using horseradish peroxidase-conjugated streptavidin. (E) HagF overexpression increases the expression of HrcA target genes in a time-dependent manner. In L2/HagF, which expresses untagged HagF from a recombinant plasmid, ATC was added at 14, 15, or 15.5 hpi. RNA was extracted at 16 hpi for qRT-PCR analysis. (F) At 37 °C, *hagF* silencing does not affect the expression of HrcA target genes. (G) At 40.5 °C, *hagF* silencing reduces the expression of the *hrcA-grpE-dnaK* operon. (F, G) ATC was added at 0 hpi, and RNA was extracted at 18 hpi.

Previous studies demonstrated that bacterial GroEL interacts with HrcA to augment its transcriptional repression activity ^22^. To determine if HagF may also interact with HrcA to regulate its transcriptional repression activity, we performed pulldown assays using purified HagF and HrcA tagged with differential epitope tags. Notably, NH-HagF pulled down STREP-tagged HrcA (NS-HrcA) (Fig. 3C); similarly, NH-HrcA also pulled down STREP-tagged HagF (NS-HagF) (Fig. 3D). These results indicate that HagF directly binds to HrcA, which may affect HrcA’s transcriptional repression activity.

To test the hypothesis that HagF functions as a regulator of HrcA, we quantified the expression of HrcA target genes in *C. trachomatis* under conditions of hagF overexpression and silencing. ATC-induced HagF overexpression in L2/HagF cultured at 37 °C led to consistent increases in the expression of *hrcA, grpE*, *dnaK*, *groES*, and *groEL* (Fig. 3E), suggesting that HagF is capable of inhibiting HrcA’s transcriptional repression activity. In contrast, ATC-induced dC12-mediated hagF silencing in both L2/dC12-hag-g1 and L2/dC12-hag-g2 did not affect the expression of any of these genes at 37 °C but significantly downregulated *hrcA, grpE,* and *dnaK* (albeit not *groES,* and groEL) at 40.5 °C (Fig. 3G). These findings indicate that HagF inhibits HrcA’s transcriptional repression of the *hrcA-grpE-dnaK* chaperone during heat shock.

### HagF inhibits HrcA’s repression activity likely by interfering with HrcA dimerization

To investigate the mechanism by which HagF inhibits the repression activity of HrcA, we employed AlphaFold3 ^41^ to predict the three-dimensional structures of both HagF and the chlamydial HrcA as well as their interactions. The predicted 3-D structure of *C. trachomatis* HagF (Fig. 4A) is similar to the experimentally determined structure of HrcA from *Thermotoga maritima* (Fig. S3) ^42^. AlphaFold Multimer predicted dimerization of the *C. trachomatis* HrcA (Fig. 4B, C), similar to the experimentally determined *T. maritima* HrcA ^42^.

**Fig. 4.**
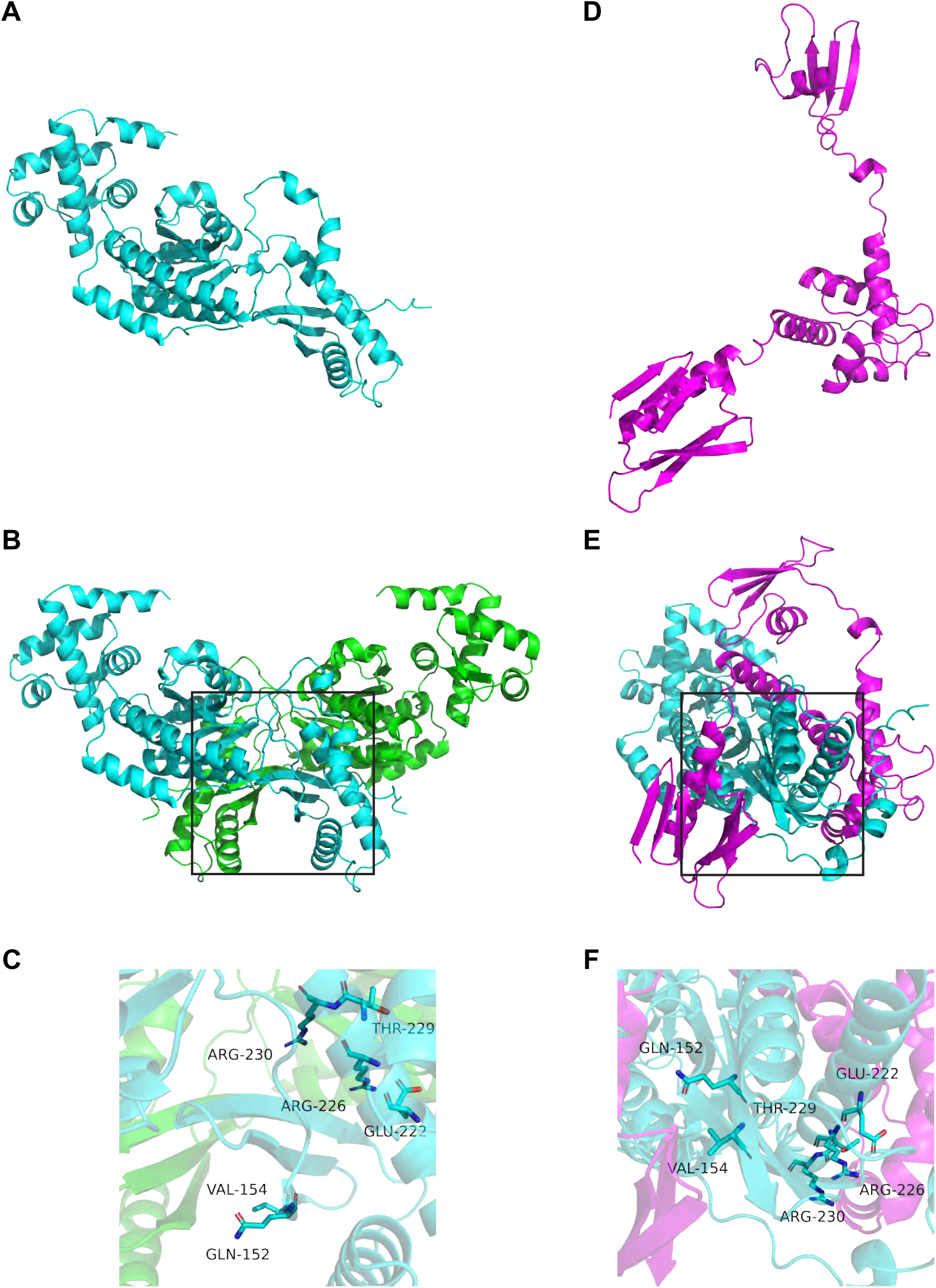
HagF inhibits HrcA homodimerization. (A) AlphaFold3-predicted tertiary structure of the HrcA monomer. (B) AlphaFold3-predicted structure of the HrcA homodimer. (C) Close-up view of residues involved in HrcA-HrcA interactions at the dimerization interface. (D) AlphaFold3-predicted tertiary structure of HagF. (E) AlphaFold3-predicted structure of the HagF-HrcA heterodimer. (F) Close-up view of interacting residues at the HrcA dimerization interface in the HagF-HrcA heterodimer, showing how HagF binding blocks homodimer formation.

There are three distinct structure domains in the AlphaFold3 model of HagF: an N-terminal domain (residues 1-73) composed of both α-helices and β-sheets, a central domain (residues 79-180) consisting of a helical bundle, and a C-terminal domain (residues 195-242) comprising both α-helices and β-sheets (Fig. 4C). The three domains, linked by flexible regions, do not interact with each other.

Consistent with the pulldown data in Fig. 3C, D, AlphaFold Multimer supported that HagF binds to HrcA (Fig. 4D, E). All three HagF domain structures are involved in HrcA binding (Fig. 4D, E). The predicted HagF-HrcA heterodimeric structure reveals that HagF occupies a key position that overlaps with the HrcA homodimerization interface (Fig. 4D, 4E). Critical residues in HrcA, such as glu152, val154, glu222, thr229, and arg230, are involved in both homodimerization (Fig. 4E) and interaction with HagF (Fig. 4D). This overlap indicates that HagF inhibits the transcriptional repression activity of HrcA by blocking HrcA homodimerization, thereby preventing it from binding to CIRCE elements and repressing the expression of its target genes.

### Unregulated HagF expression inhibits chlamydial growth at 37 °C

As shown in Fig. 3C, HagF overexpression relieves HagF repression of *grpE, dnaK, groES,* and *groEL*. To determine whether excessive HagF expression and the resulting chaperone upregulation affect chlamydial fitness under optimal growth conditions, we hypothesized that HagF overexpression-mediated increases in protein chaperones could inhibit chlamydial growth under non-stressed conditions. Supporting this hypothesis, in ATC-containing cultures, L2/CH-HagF and L2/NH-HagF formed smaller inclusions, compared ATC-free cultures (Fig. 5A), and L2/HagF exhibited significantly slower genome replication kinetics (Fig. 5B) and produced reduced progeny EBs (Fig. 5C). These findings indicate that tight repression of HagF under non-stressed conditions is critical for maintaining chlamydial growth and progeny formation.

**Fig. 5.**
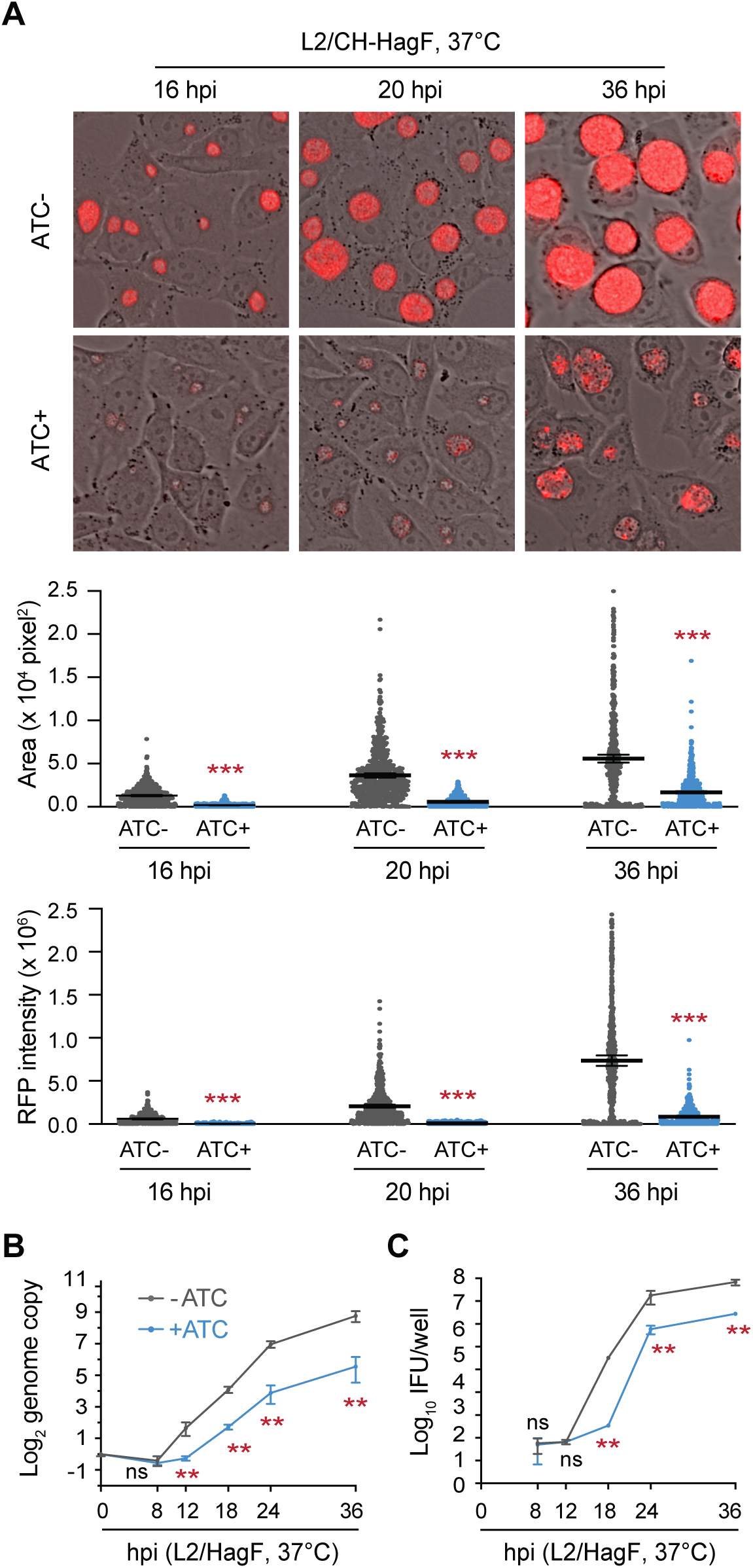
Forced HagF overexpression disrupts chlamydial growth and EB formation under non-stressed conditions. (A) ATC-induced HagF overexpression in L2/CH-HagF results in smaller inclusions with reduced red fluorescence intensity. Shown are representative fluorescence microscopy images at the indicated time points (top), with scatterplots quantifying inclusion size (middle) and RFP intensity (bottom). (B) ATC-induced HagF overexpression in L2/HagF slows chlamydial genome replication. (C) ATC-induced HagF overexpression in L2/HagF reduces progeny EB yield. (A-C) All cultures were maintained at 37 °C.

## Discussion

Bacterial survival and pathogenicity rely on precise transcriptional responses to environmental stress. One conserved regulator of this response is the transcriptional repressor HrcA, which enables many bacteria to adapt to elevated temperatures by repressing the expression of molecular chaperones. In most bacteria, HrcA binds to CIRCE elements upstream of the *hrcA-grpE-dnaK* and *groES-groEL* operons and represses the transcription of genes encoding essential chaperone proteins ^16,20,25,43-45^. While this core regulon is well established, additional HrcA-regulated genes have been identified in diverse organisms, particularly those that encounter changing environments ^16,25,44^.

In *Chlamydia*, the HrcA regulon has traditionally been thought to include only these five classical genes ^20,46^. Here, we identify *ctl0271 (hagF)* as the sixth HrcA-repressed gene. Multiple lines of evidence support this conclusion: the presence of a CIRCE-like element upstream of *hagF*, HrcA binding to its promoter in ChIP-qPCR assays, reduced *hagF* expression upon HrcA overexpression, and strong induction of *hagF* in response to heat shock. These findings expand the known HrcA regulon and suggest that the chlamydial heat shock response includes additional regulatory components beyond the core chaperone network.

While we were finalizing this study for publication, Kozusnik et al. reported the activation of *ctl0271*, the gene we designate as *hagF*, in *C. trachomatis*, and of the HagF ortholog *wcw_0502* in the chlamydia-like organism *Waddlia chondrophila*, in response to heat shock ^47^. They also proposed that these genes are targets of HrcA ^47^. However, they did not investigate the physiological function of the encoded proteins. Our findings now demonstrate that although *hagF* is dispensable for chlamydial development and growth at 37 °C, HagF becomes essential for EB formation under heat stress. This phenotype is notable given the reduced genome of *Chlamydia* and underscores the importance of adaptive regulation for survival in host environments. HagF is conserved across all sequenced members of the order Chlamydiales, including both *Chlamydia* and chlamydia-like organisms, suggesting a shared evolutionary need to couple stress response with developmental progression. While many chlamydial and host factors have been shown to promote persistence under stress, relatively few proteins are known to actively support productive development in such conditions ^9-14^. The discovery that HagF advances the productive developmental cycle during heat stress provides new insight into how Chlamydia maintains infectivity in febrile hosts.

At the molecular level, HagF binds directly to HrcA, as demonstrated by reciprocal pulldown assays. Structural modeling using AlphaFold3 supports this interaction and shows that HagF occupies the HrcA homodimerization interface, thereby preventing the formation of functional dimers required for DNA binding. All three structural domains of HagF contribute to this interaction. Importantly, *hagF* is repressed by HrcA, establishing a feedback loop that helps fine-tune chaperone gene expression under changing thermal conditions.

This reciprocal regulatory relationship between HrcA and HagF is temperature dependent and is illustrated in Fig. 6. Under non-stressed conditions, HrcA is stable and represses *hagF*, keeping HagF levels low and limiting its impact on gene expression (Fig. 6A). Previous studies suggest that during these conditions, GroEL or the GroESL complex enhances HrcA activity, likely by stabilizing the repressor and forming a broadly conserved positive feedback loop ^22-24^. This feedback mechanism helps maintain repression of chaperone genes in the absence of stress. In contrast, upon heat stress, much of the HrcA protein becomes destabilized, reducing its DNA-binding activity, although a portion remains functional (Fig. 6B). Concurrently, heat-induced *hagF* expression leads to the accumulation of HagF, which binds this residual HrcA pool, further relieving repression of chaperone genes and enhancing their expression. This HagF-mediated negative feedback loop appears to be unique to *Chlamydia* and enhances responsiveness by amplifying derepression during stress. Silencing *hagF* under these conditions reduces expression from the *hrcA-grpE-dnaK* operon but not from the *groES-groEL* operon, possibly reflecting differences in CIRCE element affinity or promoter architecture. The marked reduction in EB formation in *hagF*-silenced cells under heat stress further suggests that the DnaK-GrpE chaperone system is especially important for the secondary chlamydial differentiation in these conditions.

**Fig. 6.**
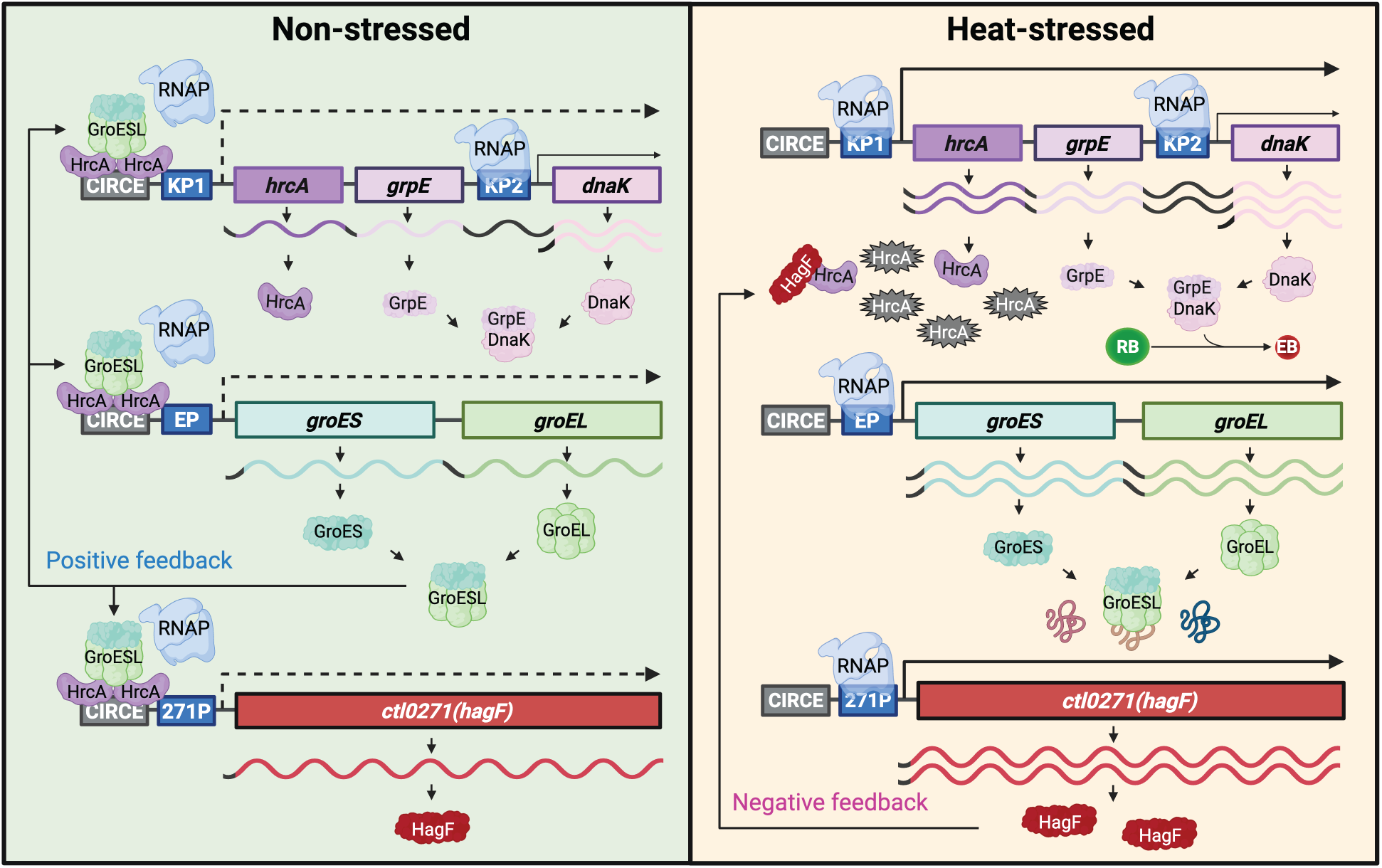
Model for dual feedback regulation of HrcA by GroESL and HagF in *Chlamydia*. Under non-stressed conditions (left), HrcA homodimers bind CIRCE elements upstream of the *hrcA-grpE-dnaK*, *groES-groEL*, and *ctl0271 (hagF)* operons, repressing transcription, as indicated by dotted lines from the KP1, EP, and 271P promoters. The GroEL-GroES (GroESL) complex enhances HrcA function, likely by stabilizing the protein, forming a broadly conserved positive feedback loop that maintains repression of chaperone genes in the absence of stress. Although not depicted, GroEL alone has also been shown to enhance HrcA activity in *Chlamydia* ^22^. Under heat-stressed conditions (right), HrcA becomes destabilized and largely inactivated, reducing its DNA-binding activity and resulting in increased transcription from the KP1, EP, and 271P promoters. Concurrently, heat-induced expression of *hagF* leads to the accumulation of HagF, which binds to the remaining HrcA protein and inhibits its dimerization. This relieves repression of chaperone operons and establishes a negative feedback loop that amplifies the heat shock response. HagF is especially important for the induction of the *hrcA-grpE-dnaK* operon and promotes progeny EB formation under stress. Heat-denatured proteins compete with HrcA for GroESL binding, disrupting the GroESL-mediated positive feedback loop. Figure was generated using a paid subscription to bioRender.

Notably, the GroEL- or GroESL-mediated positive feedback loop is broadly conserved across many bacterial species ^22-24^. In contrast, the HagF-mediated negative feedback loop appears to be unique to *Chlamydia* and its close relatives, consistent with HagF’s narrow phylogenetic distribution. This contrast underscores an evolutionary adaptation in *Chlamydia*, which has evolved a lineage-specific mechanism to regulate stress-induced gene expression and developmental progression despite lacking conventional heat shock sigma factors and operating with a reduced genome.

In summary, our study identifies *hagF* as a heat-induced, HrcA-repressed gene that encodes a direct antagonist of HrcA. This mutually inhibitory relationship integrates thermal stress signals with transcriptional and developmental control. By promoting productive development under stress, HagF enhances chlamydial fitness in febrile hosts and supports transmission to more permissive environments. Together with the observation by Kozusnik et al. that HagF orthologs are upregulated by heat shock and other stressors in environmental chlamydiae, our findings not only expand the known HrcA regulon but also reveal a *Chlamydia*-specific strategy for ensuring survival and reproductive success under adverse conditions. The HagF-mediated negative feedback loop represents a rare example of stress-induced inhibition of a transcriptional repressor and may reflect a broader paradigm in the evolution of regulatory control in minimal-genome pathogens.

## Materials and Methods

### Plasmids

Plasmids pTRL2-HagF, pTRL2-NH-HagF, and pTRL2-CH-HagF were constructed for the expression of untagged HagF, N-terminally His-tagged HagF, and C-terminally His-tagged HagF, respectively, under the control of an ATC-inducible promoter in *C. trachomatis*. All three plasmids were generated using the NEBuilder HiFi DNA Assembly Kit (New England Biolabs) as described previously ^40,48^. Plasmid backbone fragments were amplified from pASK-GFP-L2-mKate2 ^49^ or pTRL2-NHGrgA ^48^ as the template, while the HagF insert was amplified from the L2 genomic DNA.

Plasmids pTRL2-dC12-ntg, pTRL2-dC12-hagF-g1, and pTRL2-dC12-hagF-g2 were constructed for ATC-inducible dCas12 (dC12)-mediated gene silencing in *C. trachomatis*. pTRL2-dC12-ntg, which expresses a guide RNA (gRNA) with no predicted target site in *C. trachomatis*, was generated by assembling a plasmid backbone (PCR-amplified from pTRL2-His-GrgA-67m ^40^ with a CRISPR/dC12 array fragment amplified from pBbS8c-ddcpf1-rfp ^29^, a gift from Nigel S. Scrutton. pTRL2-dC12-hagF-g1 and pTRL2-dC12-hagF-g2 were derived from pTRL2-dC12-ntg by replacing the ntg sequence with gRNAs targeting hagF. The sequences of the gRNAs were 5′-CTCGGCATTGTAGTTAAGGGGTA-3′ and 5′-TATTTCCCCATACATTTCTGCTC-3′ for g1 and g2, respectively.

For recombinant protein expression in *Escherichia* coli, plasmids pET28a-NH-HrcA, pET21a-NS-HrcA, pET28a-NH-HagF, and pET21a-NS-HagF were constructed to express N-terminally His-tagged or Strep II-tagged versions of HrcA and HagF. These plasmids were derived by replacing the *grgA* coding sequence in pET28a-NH-GrgA or pET21a-NS-GrgA as described previously ^39^.

The identities of all plasmids were confirmed by Sanger or Nanopore sequencing (Quintara Biosciences, Boston, MA).

### Host cells and culture conditions

Human cervical epithelial HeLa 229 cells were purchased from ATCC. Murine fibroblast L929 cells were a kind gift from Dr. Terence S. Dermody (Vanderbilt University). Both HeLa and L929 cells were maintained as monolayers at 37 °C in 5% CO₂ using Dulbecco’s modified Eagle’s medium (DMEM) supplemented with 4.5 g/L glucose, 0.11 g/L sodium pyruvate, 20 µg/mL gentamicin, and either 10% (HeLa) or 5% (L929) fetal bovine serum (FBS) (Gibco, A52567-01).

### Chlamydiae and culture conditions

Wild-type *C. trachomatis* serovar L2 strain 434/BU (L2) was obtained from the American Type Culture Collection (ATCC) and expanded in our laboratory using L929 cells as host cells. The L2/NH-HrcA strain, which expresses N-terminally His-tagged HrcA, has been described previously ^48^. The following L2 transformants were generated by transforming wild-type L2 with ATC-inducible shuttle vectors as indicated: L2/dC12-ntg (pTRL2-dC12-ntg), L2/dC12-hagF-g1 (pTRL2-dC12-hagF-g1), L2/dC12-hagF-g2 (pTRL2-dC12-hagF-g2), L2/HagF (pTRL2-HagF), L2/NH-HagF (pTRL2-NH-HagF), and L2/CH-HagF (pTRL2-CH-HagF). Transformations and selection were performed as previously described ^40^. Clonal populations of the transformants were generated by limiting dilution ^40^. EBs were centrifuged through a 35% MD-76R gradient to remove host cell debris, further separated from RBs by centrifugation through a 40%/44%/52% MD-76R discontinuous gradient as described previously ^50^. Purified EBs were resuspended in the EB storage solution SPG (0.25 M sucrose, 10 mM sodium phosphate, 5 mM L-glutamate, pH 7.2), aliquoted, and stored at −80 °C.

### Synchronized infection for growth assays and gene expression analysis

Near-confluent HeLa monolayers cultured in 6-, 12-, or 24-well plates were infected with MD-76R gradient-purified EBs, typically at a multiplicity of infection of approximately one inclusion-forming unit (IFU) per host cell. This condition resulted in 80–90% infection efficiency, as assessed by fluorescence microscopy. To synchronize infection, inoculated monolayers were centrifuged at 900 × g and 25 °C for 20 min, then washed three times with Hank’s Balanced Salt Solution. The endpoint of the wash step was designated as 0 hour post-infection (hpi). Infected cells were then cultured in DMEM supplemented with 5% FBS and 20 µg/mL gentamicin in a humidified 37 °C incubator with 5% CO_₂_.

### Quantification of chlamydial genome copy number

Chlamydial genome copy number was used as a quantitative measure of total bacterial load, including both reticulate bodies (RBs) and elementary bodies (EBs). This method is particularly useful for assessing RB replication, since RBs are non-infectious and cannot be quantified by inclusion-forming unit (IFU) assays (see below). Genome copy number thus serves as a complementary metric to IFU-based progeny analysis.

At the indicated time points, culture medium was replaced with 0.5 mL SPG per well (12- or 24-well plates) or 1.0 mL per well (6-well plates). Infected cells were detached using Cell Lifters (Corning) and collected into 4.5 mL culture tubes. The cell suspensions were sonicated using a 130-watt ultrasonic processor equipped with a 3-mm tip, set to 35% amplitude. Sonication was applied for a total of 12 seconds in alternating 2-second on/off pulses to lyse host cells and release chlamydiae. Chlamydiae were pelleted by 10-min centrifugation at 20,000 x g and 4 °C. After carefully removing the supernatant, the pellet was resuspended in 100 µL of alkaline lysis buffer (0.1 M NaOH, 0.2 mM EDTA). Lysates were heated at 95 °C for 15 min and then neutralized by adding 400 µL of neutralization buffer (25 mM Tris-HCl, pH 7.2). The neutralized DNA samples were stored at -20 °C for subsequent analysis.

Relative chlamydial chromosome abundance was quantified by qPCR using primers targeting *grgA*. Each reaction was performed in technical duplicates using DNA samples from three biological replicates. Reactions were carried out on a QuantStudio 5 Real-Time PCR System using Power SYBR Green PCR Master Mix (Thermo Fisher Scientific, catalog #A25778).

### Inclusion-forming unit (IFU) assay

The IFU assay is functionally equivalent to a colony-forming unit assay and was used to quantify infectious EB particles in either purified EB stocks or crude cell lysates based on their ability to form inclusions in host cells. To prepare cell lysates, infected cells on multi-well plates were collected and lysed, similarly to genome copy quantification assays. Lysates were centrifuged at 500 x g for 10 min at 4 °C to remove cell debris and stored at -80 °C until further use.

EB stocks or clarified lysates from infected cultures were thawed, serially diluted 1:10 with culture medium (DMEM containing 5% fetal bovine serum, 20 µg/mL gentamicin, and 1 µg/mL cycloheximide) on 96-well plates with 2- to 3-hour-old confluent L929 cell monolayers. Plates were centrifuged at 900 x g and 25 °C for 20 min and then incubated at 37 °C for 30 hours. Since all chlamydial transformants used in this study express the far-red fluorescent protein mKate, red fluorescent inclusions were counted in live cultures using an Olympus IX-50 fluorescence microscope ^40^.

### Inclusion size and red fluorescence intensity quantification

Inclusion size and red fluorescence protein (RFP) intensity, both correlating with chlamydial growth, were quantified as previously described ^51^ with modifications. Before imaging, the culture medium was replaced with phosphate-buffered saline (PBS) containing calcium and magnesium to minimize background fluorescence. Brightfield and red fluorescence images were acquired using an Olympus IX51 fluorescence microscope with an Infinity i8-3 CMOS camera, maintaining constant exposure settings across all samples. Image overlay was performed using ACINST03 software. Inclusion size and RFP intensity were quantified using the Java-based ImageJ software.

### Chromatin immunoprecipitation (ChIP)

To prepare *C. trachomatis* cultures for ChIP, two 150-mm dishes of L929 cells were infected with L2/NH-HrcA and cultured in medium supplemented with cycloheximide (1 µg/mL final concentration). At 14 hpi, ATC was added to a final concentration of 10 nM to induce HrcA expression. At 16 hpi, crosslinking and DNA fragmentation were performed as previously described ^52^ with modifications. Medium was replaced with 15 mL of 1% formaldehyde in PBS, and crosslinking proceeded at room temperature (RT) for 20 min. Formaldehyde was then quenched by replacing it with 20 mL of quench solution (PBS containing 0.25 M glycine, pH 7.5). After incubation at RT for 15 min, the quench solution was aspirated. and cells were scraped into 15 mL of quench solution containing 0.2% bovine serum albumin (BSA), which were subject to centrifugation at 500 × g for 10 min at 4 °C.

The resulting pellet was resuspended in 1 mL of lysis buffer, made of 25 mM Tris-HCl (pH 7.5), 150 mM NaCl, 0.5% sodium deoxycholate, 1% NP-40, 0.1% SDS, and protease inhibitor cocktail c0mplete (Roche). Cells were lysed by sonication using a 130-watt Ultrasonic Processor (3-mm tip, 35% amplitude) in 5-second on/off cycles for 9 min, yielding DNA fragments between 300 and 1200 bp. The sonicated lysate was clarified by centrifugation at 6,000 × g for 10 min at 4°C and pre-cleared by incubation with 50 µL (bed volume) of Protein A/G Sepharose beads (Millipore Sigma, catalog # IP-10) for 2 h at 4 °C on a Nutator.

While pre-clearing was in progress, 5 µg of either monoclonal anti-His antibody (GenScript, A00186) or control mouse IgG (Millipore Sigma, M9269) was mixed with 80 µL of washed Protein A/G Sepharose in a 1.5-mL tube. After rotating for 1 h at 4 °C, 1 mL of 5% BSA was added, and tubes were incubated for 2 h at 4 °C to block nonspecific binding. BSA and free antibodies were removed, and 250 µL of pre-cleared lysate was added to the antibody-coated beads, followed by overnight incubation on a Nutator at 4 °C.

Beads were washed sequentially with the following buffers: low-salt wash buffer (20 mM Tris-HCl pH 8.1, 150 mM NaCl, 2 mM EDTA, 1% Triton X-100, 0.1% SDS), high-salt wash buffer (same as above, but with 500 mM NaCl), LiCl wash buffer (10 mM Tris-HCl pH 8.1, 0.25 M LiCl, 1 mM EDTA, 1% NP-40, 1% sodium deoxycholate), and twice with TE buffer (10 mM Tris-HCl pH 8.1, 1 mM EDTA).

Protein–DNA complexes were eluted using 200 µL of freshly prepared elution buffer (1% SDS, 0.1 M NaHCO₃). After brief centrifugation, supernatants were transferred to new tubes. To reverse crosslinking, 8 µL of 5 M NaCl and 4 µL of RNase A (10 mg/mL) were added, and tubes were incubated overnight at 65 °C with shaking. Protein digestion was performed by adding 2 µL of proteinase K (20 mg/mL), 16 µL of 0.5 M EDTA (pH 8.0), and 8 µL of 1 M Tris (pH 6.5), followed by incubation at 45 °C for 2 h with shaking. ChIP DNA was purified using the QIAquick PCR Purification Kit (Qiagen) and stored at -20 °C until further use.

### Quantification of gene promoters in ChIP DNA

Enrichment of target gene promoters in ChIP DNA was assessed by qPCR using DNA precipitated with anti-His antibody or control IgG and a non-precipitated input DNA control. PCR reactions were performed using promoter-specific primers (Table S1) as described for genome copy number qualification. Promoter enrichment was calculated relative to input DNA and compared between anti-His and IgG ChIP to assess HrcA binding specificity.

### RNA extraction and purification

*C. trachomatis*-infected cells cultured in 6- or 24-well plates were lysed directly in TRI reagent (Millipore Sigma, catalog no. 93289). Total RNA was extracted from the aqueous phase according to the manufacturer’s instructions and treated with two successive rounds of DNase I-XT digestion (New England Biolabs) to remove residual genomic DNA. Complete DNA removal was verified by PCR analysis ^40^. RNA concentration and purity were determined using Qubit RNA assay kits (Thermo Fisher Scientific). DNA-free RNA was aliquoted and stored at -80 °C until use.

### Quantification of gene expression

We performed quantitative reverse transcription-PCR (qRT-PCR) using a QuantStudio 5 Real-Time PCR System (Thermo Fisher Scientific) and the Luna Universal One-Step qRT-PCR Kit (New England BioLabs, catalog #E3005E) to quantify gene expression in *Chlamydia trachomatis* as described previously ^40,48^. RNA samples stored at –80 °C were thawed on ice and diluted 1:200 in nuclease-free water. Each 10 µL reaction contained 5 µL of Luna Universal One-Step Reaction Mix, 0.5 µL of 20X Luna WarmStart Reaction Enzyme Mix, 0.4 µL each of 10 µM forward and reverse primers, and 3.7 µL of diluted RNA. In addition to assays for target genes (*hagF*, *hrcA*, *grpE*, *dnaK*, *groES*, *groEL*, *dC12*), qRT-PCR was also performed for 23S rRNA, which served as the internal reference. We selected 23S rRNA because our previous studies showed that its expression is unaffected by heat shock or short-term (2-h) *hrcA* overexpression ^5^. Relative gene expression levels were calculated using the 2^–ΔΔCT method ^53^. All experiments were conducted using three or five biologically independent RNA samples, each measured in technical duplicate. Information for qRT-PCR primers is provided in Table S1.

### Expression of HIS- and Strep II-tagged proteins in and their purification from *E. coli*

*E. coli* BL21 (DE3) pLysS cells were transformed with plasmids pET28a-NH-HrcA, pET21a-NS-HrcA, pET28a-NH-HagF, or pET21a-NS-HagF. Protein expression was induced as previously described ^39^. His-tagged proteins (NH-HrcA and NH-HagF) were purified using HisPur cobalt Sepharose (Thermo Fisher Scientific) following established protocols ^39^. Strep II-tagged proteins (NS-HrcA and NS-HagF) were purified using Strep-Tactin XT Sepharose according to the manufacturer’s instructions.

### Protein pulldown assays

Pulldown assays were performed as previously described with modifications ^54^. For each pulldown reaction, 20 µg of purified NH-HagF or NH-HrcA was incubated with 20 µg of recombinant NS-HrcA or NS-HagF, respectively, in a total volume of 40 µL. In negative control reactions, NH-HagF or NH-HrcA was replaced with protein storage buffer (100 mM NaCl, 50 mM sodium phosphate, pH 7.0). Protein mixtures were incubated at 37 °C for 30 min. During the incubation, 50 µL of HisPur cobalt Sepharose suspension (Thermo Fisher Scientific) was washed twice with 1 mL of HisPur purification wash buffer containing 50 mM imidazole, followed by two washes with 200 µL of wash buffer containing 10 mM imidazole. Protein mixtures were then added to the imidazole-prewashed beads and incubated on a Nutator at 4 °C for 1 h. After binding, beads were washed five times with 1 mL of wash buffer containing 10 mM imidazole and 1% NP-40, followed by two washes with 1 mL of wash buffer containing 50 mM imidazole. Bound proteins were eluted in 30 µL of elution buffer (50 mM sodium phosphate, pH 7.4, 300 mM NaCl, 150 mM imidazole). Eluted proteins were resolved by 12% SDS-PAGE and transferred to a PVDF membrane. Membranes were blocked with 3% BSA and probed with horseradish peroxidase-conjugated streptavidin. Signal was detected by chemiluminescence (Millipore Sigma, catalog # OR03L).

### Immunofluorescence assays

cells cultured in 6-well plates were infected with L2/NH-HagF or L2/CH-HagF and incubated at 37 °C. At 16 hpi, anhydrotetracycline (ATC) was added to a final concentration of 10 nM to induce NH-HagF or CH-HagF expression. At 22 hpi, cells were fixed sequentially with 4% paraformaldehyde followed by cold methanol. NH-HagF and CH-HagF were detected using a mouse anti-His antibody (Abcam, ab9108) and a TRITC-conjugated goat anti-mouse IgG secondary antibody (Millipore Sigma, catalog #T5393). GrgA was visualized using a rabbit anti-GrgA antibody ^48^ and a FITC-conjugated goat anti-rabbit IgG (Immunotech, catalog # IM0833). Fluorescence images were acquired using an Infinity i8-3 CMOS monochrome camera. Image overlay and color merging were performed using ACINST03 software ^40^.

### Protein 3-dimensional structure Prediction

The three-dimensional structures of HagF, HrcA, the HrcA homodimer, and the HagF-HrcA heterodimer were predicted using the AlphaFold3 server ^41^. Structure predictions were based on full-length amino acid sequences, and multimeric models were generated using AlphaFold’s multimer mode where applicable.

### Statistical analysis

Statistical comparisons between two groups were performed using two-tailed *t*-tests. Where applicable, *P* values were adjusted for multiple comparisons using the Benjamini-Hochberg procedure to control the false discovery rate. Comparisons involving three or more groups were conducted using one-way ANOVA followed by Dunnett’s post hoc tests.

## Acknowledgments

This work was supported by the National Institutes of Health grant numbers AI140167 and AI154305 (to H.F.) and AI182210 (to G.Z. and H.F.). Y.H. was supported by a scholarship (award no. 201806370171) from the China Scholarship Council (CSC) from September 2018 to September 2020.

## References

1 Roncarati, D., Vannini, A. & Scarlato, V. Temperature sensing and virulence regulation in pathogenic bacteria. Trends Microbiol (2024). 10.1016/j.tim.2024.07.009

2 Balchin, D., Hayer-Hartl, M. & Hartl, F. U. Recent advances in understanding catalysis of protein folding by molecular chaperones. FEBS Lett 594, 2770–2781 (2020). 10.1002/1873-3468.13844

3 Sharp, J. D. et al. Comprehensive Definition of the SigH Regulon of Mycobacterium tuberculosis Reveals Transcriptional Control of Diverse Stress Responses. PLoS One 11, e0152145 (2016). 10.1371/journal.pone.0152145

4 Schumann, W. Regulation of bacterial heat shock stimulons. Cell Stress Chaperones 21, 959–968 (2016). 10.1007/s12192-016-0727-z

5 Huang, Y. et al. Robust heat shock response in *Chlamydia* lacking a typical heat shock sigma factor. Front Microbiol 12, 812448 (2021). 10.3389/fmicb.2021.812448

6 Elwell, C., Mirrashidi, K. & Engel, J. Chlamydia cell biology and pathogenesis. Nat Rev Microbiol 14, 385–400 (2016). 10.1038/nrmicro.2016.30

7 Hybiske, K. & Stephens, R. S. Mechanisms of *Chlamydia trachomatis* entry into nonphagocytic cells. Infect Immun 75, 3925–3934 (2007). 10.1128/iai.00106-07

8 Hybiske, K. & Stephens, R. S. Mechanisms of host cell exit by the intracellular bacterium *Chlamydia*. Proc Natl Acad Sci USA 104, 11430–11435 (2007). 10.1073/pnas.0703218104

9 Rockey, D. D., Wang, X., Debrine, A., Grieshaber, N. & Grieshaber, S. S. Metabolic dormancy in Chlamydia trachomatis treated with different antibiotics. Infect Immun 92, e0033923 (2024). 10.1128/iai.00339-23

10 Pokorzynski, N. D., Alla, M. R. & Carabeo, R. A. Host Cell Amplification of Nutritional Stress Contributes To Persistence in Chlamydia trachomatis. mBio, e0271922 (2022). 10.1128/mbio.02719-22

11 Shima, K. et al. Regulation of the mitochondrion-fatty acid axis for the metabolic reprogramming of *Chlamydia trachomatis* during treatment with beta-lactam antimicrobials. mBio 12 (2021). 10.1128/mBio.00023-21

12 Brockett, M. R. & Liechti, G. W. Persistence Alters the Interaction between Chlamydia trachomatis and Its Host Cell. Infect Immun 89, e0068520 (2021). 10.1128/iai.00685-20

13 Huston, W. M., Theodoropoulos, C., Mathews, S. A. & Timms, P. Chlamydia trachomatis responds to heat shock, penicillin induced persistence, and IFN-gamma persistence by altering levels of the extracytoplasmic stress response protease HtrA. BMC Microbiol 8, 190 (2008). 10.1186/1471-2180-8-190

14 Schuchardt, L. & Rupp, J. Chlamydia trachomatis as the Cause of Infectious Infertility: Acute, Repetitive or Persistent Long-Term Infection? Curr Top Microbiol Immunol 412, 159–182 (2018). 10.1007/82_2016_15

15 Panzetta, M. E., Valdivia, R. H. & Saka, H. A. Chlamydia Persistence: A Survival Strategy to Evade Antimicrobial Effects in-vitro and in-vivo. Front Microbiol 9 (2018). 10.3389/fmicb.2018.03101

16 Holmes, C. W., Penn, C. W. & Lund, P. A. The hrcA and hspR regulons of *Campylobacter jejuni*. Microbiology 156, 158–166 (2010). 10.1099/mic.0.031708-0

17 Spohn, G. et al. Dual control of *Helicobacter pylori* heat shock gene transcription by HspR and HrcA. J Bacteriol 186, 2956–2965 (2004). 10.1128/jb.186.10.2956-2965.2004

18 Lund, P. A. Microbial molecular chaperones. Adv Microb Physiol 44, 93–140 (2001). 10.1016/s0065-2911(01)44012-4

19 Baldini, R. L., Avedissian, M. & Gomes, S. L. The CIRCE element and its putative repressor control cell cycle expression of the *Caulobacter crescentus groESL* operon. J Bacteriol 180, 1632–1641 (1998). 10.1128/jb.180.7.1632-1641.1998

20 Wilson, A. C. & Tan, M. Stress response gene regulation in Chlamydia is dependent on HrcA-CIRCE interactions. J Bacteriol 186, 3384–3391 (2004).

21 Wilson, A. C. & Tan, M. Functional analysis of the heat shock regulator HrcA of Chlamydia trachomatis. J Bacteriol 184, 6566–6571 (2002).

22 Wilson, A. C., Wu, C. C., Yates, J. R., 3rd & Tan, M. Chlamydial GroEL autoregulates its own expression through direct interactions with the HrcA repressor protein. J Bacteriol 187, 7535–7542 (2005).

23 Mogk, A. et al. The GroE chaperonin machine is a major modulator of the CIRCE heat shock regulon of Bacillus subtilis. Embo j 16, 4579–4590 (1997). 10.1093/emboj/16.15.4579

24 Grandvalet, C., Rapoport, G. & Mazodier, P. hrcA, encoding the repressor of the groEL genes in Streptomyces albus G, is associated with a second dnaJ gene. J Bacteriol 180, 5129–5134 (1998). 10.1128/jb.180.19.5129-5134.1998

25 Hecker, M., Schumann, W. & Volker, U. Heat-shock and general stress response in Bacillus subtilis. Mol Microbiol 19, 417–428 (1996). 10.1046/j.1365-2958.1996.396932.x

26 Rusanganwa, E. & Gupta, R. S. Cloning and characterization of multiple groEL chaperonin-encoding genes in Rhizobium meliloti. Gene 126, 67–75 (1993). 10.1016/0378-1119(93)90591-P

27 Roncarati, D. & Scarlato, V. Regulation of heat-shock genes in bacteria: from signal sensing to gene expression output. FEMS Microbiol Rev 41, 549–574 (2017). 10.1093/femsre/fux015

28 Abdeen, S. et al. GroEL/ES inhibitors as potential antibiotics. Bioorganic & medicinal chemistry letters 26, 3127–3134 (2016). 10.1016/j.bmcl.2016.04.089

29 Jervis, A. J. et al. A plasmid toolset for CRISPR-mediated genome editing and CRISPRi gene regulation in *Escherichia coli*. Microbial Biotechnology 14, 1120–1129 (2021). 10.1111/1751-7915.13780

30 da Cunha, M. et al. Identification of type III secretion substrates of Chlamydia trachomatis using Yersinia enterocolitica as a heterologous system. BMC Microbiol 14, 40 (2014). 10.1186/1471-2180-14-40

31 Steiert, B. et al. Global mapping of the Chlamydia trachomatis conventional secreted effector-host interactome reveals CebN interacts with nucleoporins and Rae1 to impede STAT1 nuclear translocation. bioRxiv (2024). 10.1101/2024.04.25.587017

32 Cambronne, E. D. & Roy, C. R. Recognition and delivery of effector proteins into eukaryotic cells by bacterial secretion systems. Traffic 7, 929–939 (2006).

33 Hueck, C. J. Type III protein secretion systems in bacterial pathogens of animals and plants. Microbiol Mol Biol Rev 62, 379–433 (1998).

34 Wagner, S. et al. Bacterial type III secretion systems: a complex device for the delivery of bacterial effector proteins into eukaryotic host cells. FEMS Microbiol Lett 365 (2018). 10.1093/femsle/fny201

35 Voth, D. E., Broederdorf, L. J. & Graham, J. G. Bacterial Type IV secretion systems: versatile virulence machines. Future Microbiol 7, 241–257 (2012). 10.2217/fmb.11.150

36 Freudl, R. Signal peptides for recombinant protein secretion in bacterial expression systems. Microb Cell Fact 17, 52 (2018). 10.1186/s12934-018-0901-3

37 Angot, A., Vergunst, A., Genin, S. & Peeters, N. Exploitation of eukaryotic ubiquitin signaling pathways by effectors translocated by bacterial type III and type IV secretion systems. PLoS Pathog 3, e3 (2007). 10.1371/journal.ppat.0030003

38 Wang, J., Brodmann, M. & Basler, M. Assembly and Subcellular Localization of Bacterial Type VI Secretion Systems. Annu Rev Microbiol 73, 621–638 (2019). 10.1146/annurev-micro-020518-115420

39 Bao, X., Nickels, B. E. & Fan, H. *Chlamydia trachomatis* protein GrgA activates transcription by contacting the nonconserved region of σ^66^. Proc Natl Acad Sci USA 109, 16870–16875 (2012). 10.1073/pnas.1207300109

40 Lu, B. et al. Requirement of GrgA for *Chlamydia* infectious progeny production, optimal growth, and efficient plasmid maintenance. mBio 15, e0203623 (2024). 10.1128/mbio.02036-23

41 Abramson, J. et al. Accurate structure prediction of biomolecular interactions with AlphaFold 3. Nature 630, 493–500 (2024). 10.1038/s41586-024-07487-w

42 Liu, J. et al. Crystal structure of a heat-inducible transcriptional repressor HrcA from Thermotoga maritima: structural insight into DNA binding and dimerization. J Mol Biol 350, 987–996 (2005). 10.1016/j.jmb.2005.04.021

43 de Barsy, M. et al. Regulatory (pan-)genome of an obligate intracellular pathogen in the PVC superphylum. The ISME Journal 10, 2129–2144 (2016). 10.1038/ismej.2016.23

44 Roncarati, D. et al. Helicobacter pylori Stress-Response: Definition of the HrcA Regulon. Microorganisms 7 (2019). 10.3390/microorganisms7100436

45 Hanson, B. R. & Tan, M. Transcriptional regulation of the *Chlamydia* heat shock stress response in an intracellular infection. Mol Microbiol 97, 1158–1167 (2015). 10.1111/mmi.13093

46 Tan, M., Wong, B. & Engel, J. N. Transcriptional organization and regulation of the dnaK and groE operons of Chlamydia trachomatis. J Bacteriol 178, 6983–6990 (1996).

47 Kozusnik, T., Kebbi-Beghdadi, C., Ardissone, S., Adams, S. E. & Greub, G. A conserved Chlamydiota-specific Type III Secretion System effector linked to stress response. Microbiology (Reading) 171 (2025). 10.1099/mic.0.001545

48 Wurihan, W. et al. Identification of a GrgA-Euo-HrcA transcriptional regulatory network in *Chlamydia*. mSystems 6, e0073821 (2021). 10.1128/mSystems.00738-21

49 Wickstrum, J., Sammons, L. R., Restivo, K. N. & Hefty, P. S. Conditional gene expression in *Chlamydia trachomatis* using the tet system. PLoS One 8, e76743 (2013). 10.1371/journal.pone.0076743

50 Wan, D. et al. Analyzing RNA-Seq data from Chlamydia with super broad transcriptomic activation: challenges, solutions, and implications for other systems. BMC Genomics 25, 801 (2024). 10.1186/s12864-024-10714-3

51 Wurihan, W. et al. GrgA overexpression inhibits *Chlamydia trachomatis* growth through sigma66- and sigma28-dependent mechanisms. Microb Pathog 156, 104917 (2021). doi10.1016/j.micpath.2021.104917

52 Hanson, B. R. & Tan, M. Intra-ChIP: studying gene regulation in an intracellular pathogen. Curr Genet (2016). 10.1007/s00294-016-0580-8

53 Livak, K. J. & Schmittgen, T. D. Analysis of relative gene expression data using real-time quantitative PCR and the 2(-Delta Delta C(T)) Method. Methods 25, 402–408 (2001). 10.1006/meth.2001.1262

54 Desai, M. et al. Role for GrgA in regulation of sigma(28)-dependent transcription in the obligate intracellular bacterial pathogen *Chlamydia trachomatis*. J Bacteriol 200 (2018). 10.1128/JB.00298-18

