## Supplementary material for "A lineage-specific heat-induced feedback loop controls HrcA to promote chlamydial fitness under stress": Fig. S1

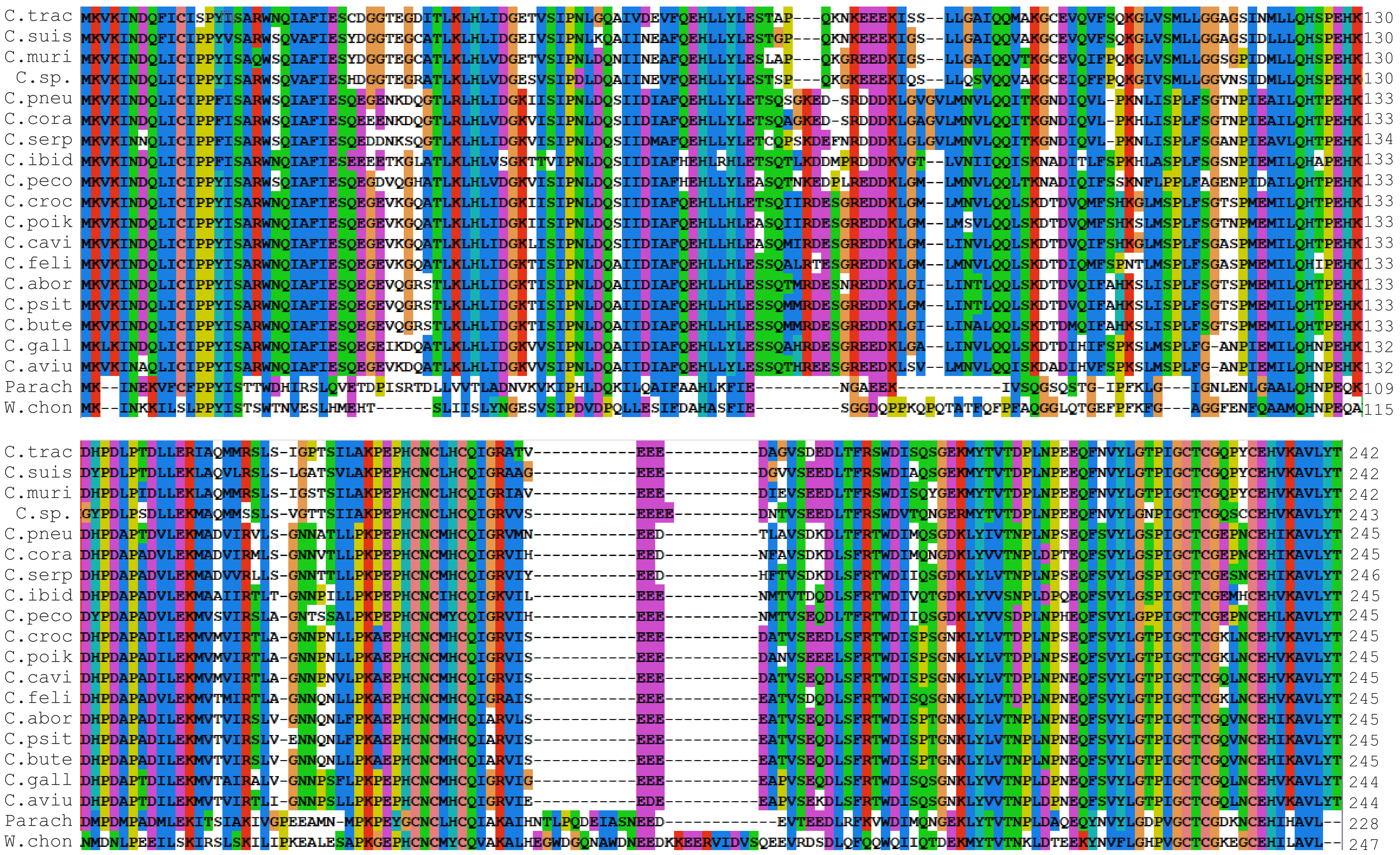

Fig. S1. Sequence alignment of *hagF* orthologs from *Chlamydia* species and selected *Chlamydia*-like organisms.
