## Supplementary material for "A lineage-specific heat-induced feedback loop controls HrcA to promote chlamydial fitness under stress": Fig. S2

**A**

```
C.trachomatis HagF          1   MKVKINDQFICISPYISARWNQIAFIESCDGGTEGDITLKLHLIDGETVSIPNLGQAIVD   60
                                MK+KINDQ ICI PYISARWNQIAFIES +G  +   TLKLHLIDG+ VSIPNL QAI+D
B.fragilis S6L5 EYE60928.1  1   MKLKINDQLICIPPYISARWNQIAFIESQEGEIKDQATLKLHLIDGKVVSIPNLDQAIID   60

C.trachomatis HagF          61  EVFQEHLLYLESTAPQKN---KEEEKISSLLGAIQQMAKGCEVQVFSQKGLVSMLLGGAG   117
                                  FQEHLLYLES+   ++   +EE+K+ +L+  +QQ++K  ++ +FS K L+S L G A
B.fragilis S6L5 EYE60928.1  61  IAFQEHLLYLESSQAHRDESGREEDKLGALINVLQQLSKDTDIHIFSPKSLMSPLFG-AN   119

C.trachomatis HagF          118 SINMLLQHSPEHKDHPDLPTDLLERIAQMMRSLSIGPTSILAKPEPHCNCLHCQIGRATV   177
                                 I M+LQH+PEHKDHPD PTD+LE++   +R+L     S L KPEPHCNC+HCQIGR
B.fragilis S6L5 EYE60928.1  120 PIEMILQHNPEHKDHPDAPTDILEKMVTAIRALVGNNPSFLPKPEPHCNCMHCQIGRVIG   179

C.trachomatis HagF          178 EEEDAGVSDEDLTFRSWDISQSGEKMYTVTDPLNPEEQFNVVYLGTPIGCTCGQPYCEHVK   237
                                EEE+A VS++DL+FR+WDISQSG K+Y VT+PL+P EQF+VVYLGTPIGCTCGQ  CEHVK
B.fragilis S6L5 EYE60928.1  180 EEEEAPVSEQDLSFRTWDISQSGNKLYVVTNPLDPNEQFSVVYLGTPIGCTCGQLNCEHVK   239

C.trachomatis HagF          238 AVLYT 242
                                AVLYT
B.fragilis S6L5 EYE60928.1  240 AVLYT 244
```

**B**

```
B.fragilis S6L5 EYE60928.1   1   IAFQEHLLYLESSQAHRDESGREEDKLGALINVLQQLSKDTDIHIFSPKSLMSPLFGANP   60
                                 IAFQEHLLYLESSQAHRDESGREEDKLGALINVLQQLSKDTDIHIFSPKSLMSPLFGANP
C.gallinacea WP_021828457.1  1   IAFQEHLLYLESSQAHRDESGREEDKLGALINVLQQLSKDTDIHIFSPKSLMSPLFGANP   60

B.fragilis S6L5 EYE60928.1   61  IAFQEHLLYLESSQAHRDESGREEDKLGALINVLQQLSKDTDIHIFSPKSLMSPLFGANP   120
                                 IAFQEHLLYLESSQAHRDESGREEDKLGALINVLQQLSKDTDIHIFSPKSLMSPLFGANP
C.gallinacea WP_021828457.1  61  IAFQEHLLYLESSQAHRDESGREEDKLGALINVLQQLSKDTDIHIFSPKSLMSPLFGANP   120

B.fragilis S6L5 EYE60928.1   121 IEMILQHNPEHKDHPDAPTDILEKMVTAIRALVGNNPSFLPKPEPHCNCMHCQIGRVIGE   180
                                 IEMILQHNPEHKDHPDAPTDILEKMVTAIRALVGNNPSFLPKPEPHCNCMHCQIGRVIGE
C.gallinacea WP_021828457.1  121 IEMILQHNPEHKDHPDAPTDILEKMVTAIRALVGNNPSFLPKPEPHCNCMHCQIGRVIGE   180

B.fragilis S6L5 EYE60928.1   181 EEEAPVSEQDLSFRTWDISQSGNKLYVVVTNPLDPNEQFSVVYLGTPIGCTCGQLNCEHVKA   240
                                 EEEAPVSEQDLSFRTWDISQSGNKLYVVVTNPLDPNEQFSVVYLGTPIGCTCGQLNCEHVKA
C.gallinacea WP_021828457.1  181 EEEAPVSEQDLSFRTWDISQSGNKLYVVVTNPLDPNEQFSVVYLGTPIGCTCGQLNCEHVKA   240

B.fragilis S6L5 EYE60928.1   241 VLYT 244
                                 VLYT
C.gallinacea WP_021828457.1  241 VLYT 244
```

Fig. S2. *Bacteroides fragilis* S6L5 does not encode hrcA. (A) Pairwise sequence alignment shows a high degree of identity between CTL0271 (HagF) and a hypothetical protein from *B. fragilis* S6L5 retrieved from NCBI. (B) Pairwise alignment reveals that the *B. fragilis* S6L5 protein is identical to the HagF ortholog from *Chlamydia gallinacea.*
