## Supplementary material for "A lineage-specific heat-induced feedback loop controls HrcA to promote chlamydial fitness under stress": Fig. S3

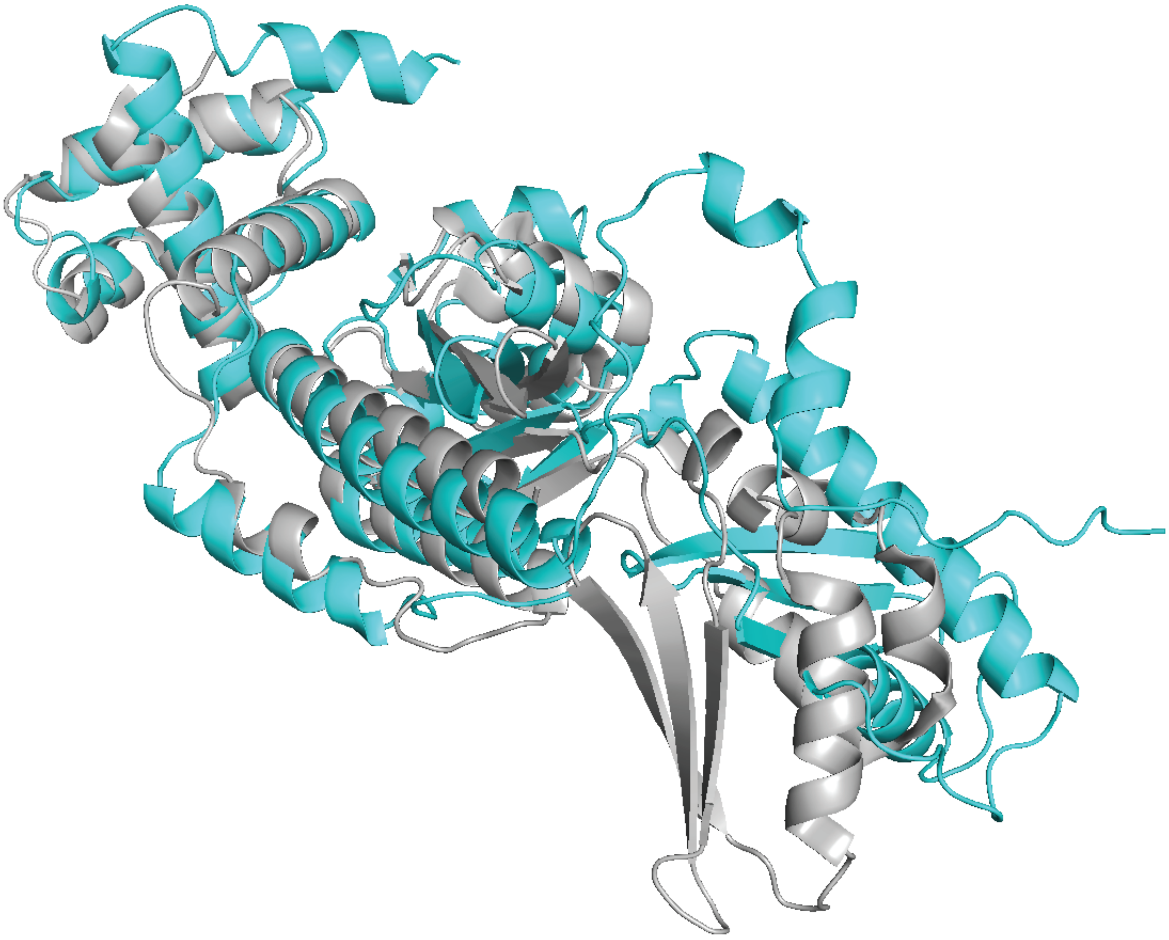

Fig. S3. Structural alignment of the AlphaFold-predicted model of C. trachomatis HrcA with the experimentally determined Thermotoga maritima HrcA structure (1STZ). The root mean square deviation (RMSD) of the alignment is 2.6 Å.
